# *brca2*-mutant zebrafish exhibit context- and tissue-dependent alterations in cell phenotypes and response to injury

**DOI:** 10.1101/2021.08.20.457131

**Authors:** Vassili A. Kouprianov, Aubrie A. Selmek, Jordan L. Ferguson, Xiaokui Mo, Heather R. Shive

**Affiliations:** Altis Biosystems, Durham, North Carolina, 27709, USA; The Ohio State University College of Veterinary Medicine, Department of Veterinary Biosciences, Columbus, Ohio, 43210, USA; North Carolina State University College of Veterinary Medicine, Department of Molecular Biomedical Sciences, Raleigh North Carolina, 27601, USA; The Ohio State University Department of Biomedical Informatics, Columbus, Ohio, 43210

## Abstract

Cancer cells frequently co-opt molecular programs that are normally activated in specific contexts, such as embryonic development and the response to injury. Determining the impact of cancer-associated mutations on cellular phenotypes within these discrete contexts can provide new insight into how such mutations lead to dysregulated cell behaviors and subsequent cancer onset. Here we assess the impact of heritable *BRCA2* mutation on embryonic development and the injury response using a zebrafish model (*Danio rerio*). Unlike most mouse models for *BRCA2* mutation, *brca2*-mutant zebrafish are fully viable and thus provide a unique tool for assessing both embryonic and adult phenotypes. We find that maternally provided brca2 is critical for normal oocyte development and embryonic survival in zebrafish, suggesting that embryonic lethality associated with BRCA2 mutation is likely to reflect defects in both meiotic and embryonic developmental programs. On the other hand, we find that adult *brca2*-mutant zebrafish exhibit aberrant proliferation of several cell types under basal conditions and in response to injury in tissues at high risk for cancer development. These divergent effects exemplify the often-paradoxical outcomes that occur in embryos (embryonic lethality) versus adult animals (cancer predisposition) with mutations in cancer susceptibility genes such as BRCA2. The altered cell behaviors identified in *brca2*-mutant embryonic and adult tissues, particularly in adult tissues at high risk for cancer, indicate that the effects of BRCA2 mutation on cellular phenotypes are both context- and tissue-dependent.

## Introduction

Carcinogenesis depends on redeployment, misuse, and dysregulation of numerous normal molecular and cellular programs. These normal programs also regulate two fundamentally important processes: embryonic development and inflammatory/injury responses. Both processes involve basic cell behaviors (e.g., cell proliferation and migration) and more complex multi-cellular processes (e.g., angiogenesis and stromal/tissue remodeling) that are misappropriated during cancer initiation and progression (reviewed in ^1–5^).

Cancer-associated genetic alterations can modify both inflammatory/injury-associated and developmental processes. Tumors are often described as “wounds that do not heal”^6^, and there is compelling evidence that chronic injury and inflammation can cause cancer^2^. Multiple tissues exhibit an intimate and synergistic association between heritable or somatically acquired genetic mutations, cellular injury and inflammation, and cancer predisposition^7–10^. On the other hand, mutation or loss of cancer-associated genes can induce developmental effects in embryos that differ significantly from effects in adult animals. This frequently manifests as paradoxical embryonic lethality versus adult cancer susceptibility; examples include well-known tumor suppressor genes (Pten^11,12^, Rb^13–15^), mediators of the DNA damage response (Atr^16,17^, Rad51^18,19^, Chek1^20^), and others^21,22^. The constraints imposed by early embryonic lethality in these models is a limitation for investigations into how and why embryonic and adult cell populations respond so differently to certain cancer-associated genetic mutations.

The tumor suppressor gene BRCA2 exemplifies this conundrum: while BRCA2 mutation is associated with cancer susceptibility in humans and animals, mouse models with homozygous Brca2 mutation exhibit early embryonic lethality^23,24^. The zebrafish (*Danio rerio*) is a freshwater fish species that provides an excellent complement to traditional mouse models for comparative cancer research. We have previously described a *brca2*-mutant zebrafish model in which the *brca2^Q658X^* mutation (nonsense mutation; RAD51 binding domain) is similar in location and type to pathologic *BRCA2* mutations associated with human cancer^25^. The resultant truncated protein lacks the majority of domains required for BRCA2 function. Despite this, *brca2* homozygous zebrafish derived from heterozygous parents are fully viable and survive to adulthood. Since cancer susceptibility in zebrafish with *brca2* mutation alone is low^25^, we use zebrafish with combined mutations in *brca2* and *tp53^26^* for carcinogenesis studies^25,27–29^. The *brca2*-mutant zebrafish model provides a unique *in vivo* system for determining how loss of functional BRCA2 affects various developmental, adult, and cancer-associated phenotypes^25,27–29^.

In the current investigation, we used our zebrafish model to further define the role for BRCA2 in embryogenesis and to determine how BRCA2 mutation affects adult cell phenotypes in the context of tissue injury/inflammation. The response to injury in zebrafish is distinguished by robust regenerative capacity in multiple adult tissues, including heart, tail fin, retina and optic nerve, and others^30,31^. We focused on the optic nerve pathway (ONP) to evaluate the relationship between BRCA2 mutation, injury response, and cancer risk because we recently reported that *brca2*-mutant/*tp53*-mutant zebrafish are at high risk for cancers in this site^27^. Interestingly, the ONP in adult fish also exhibits several characteristics that promote nerve regeneration and remyelination after injury ^32–35^. Included among these is a permissive microenvironment in which multiple non-neuronal cell populations, including astrocytes, oligodendrocytes, and local inflammatory cells, support axonal sprouting and regrowth to enable optic nerve regeneration^32,33,36–38^. We thus speculated that these features of the ONP might contribute to increased potential for tumorigenesis.

Our investigations revealed that although *brca2* homozygous embryos derived from heterozygous mutant parents are fully viable and survive to adulthood, embryos lacking maternally provided brca2 exhibit profound proliferation arrest and embryonic lethality. We further determined that oocytes from *brca2*-mutant females exhibit abnormal nuclear morphology, suggesting that brca2-associated disruptions during meiosis contribute to embryonic developmental defects. In adult zebrafish, we identified aberrant proliferative responses associated with *brca2* mutation in the cancer-prone ONP in both unperturbed and post-injury states. This includes the identification of a putative precancerous population that is highly prevalent in *brca2*-mutant/*tp53*-mutant zebrafish. Finally, we show that precancerous and cancerous lesions affecting the ONP occur at high prevalence in *brca2*-mutant/*tp53*-mutant zebrafish and are frequently bilateral. This unique vertebrate model thus allows us to identify BRCA2-associated phenotypes that are influenced by temporal, contextual, and tissue-specific factors.

## Results

### Proliferative and developmental defects occur in zebrafish embryos and oocytes lacking brca2

We have previously shown that zebrafish embryos receive abundant maternal RNA for *brca2*, which is present at the two-cell stage and persists to at least the onset of zygotic gene activation^25^. As a result, the effects of *brca2* loss during early zebrafish embryogenesis are not captured in *brca2 m/m* zebrafish embryos derived from incrosses of *brca2 +/m* parents. Adult *brca2* homozygotes cannot be used for breeding because they develop exclusively as sterile males, reflecting the requirement for *brca2* in spermatogenesis^25^ and the influence of germ cell survival on zebrafish sex differentiation^39–41^. However, concomittant homozygous mutations in *brca2* and *tp53* (*tp53^M214K^* mutation; missense mutation in p53 DNA binding domain^26^) rescue female development^25^. We therefore outcrossed *brca2 m/m;tp53 m/m* female zebrafish to wild type males in order to generate embryos lacking maternal RNA for *brca2* (**Fig. 1**). Similarly, *tp53 m/m* female zebrafish were outcrossed to wild type males (**Fig. 1**); the *tp53 m/m* zebrafish line exhibits normal fertility^26^.

**Figure 1.**
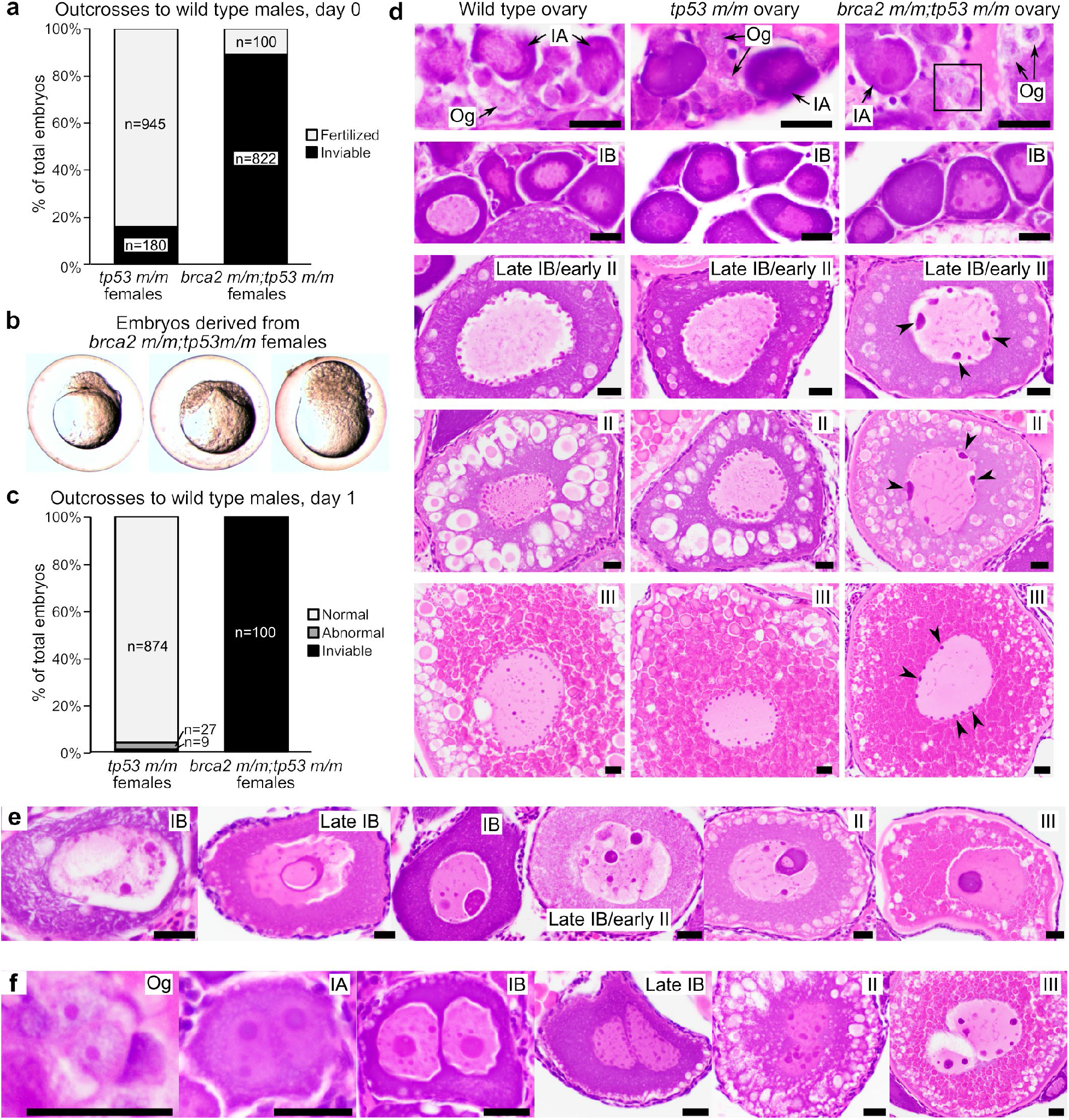
Embryos from *brca2 m/m;tp53 m/m* female zebrafish undergo proliferation arrest and death and oocytes exhibit abnormal nuclear morphology. **(a)** A minority of embryos derived from *brca2 m/m;tp53 m/m* females are viable and fertilized compared to *tp53 m/m* females. Females were outcrossed to fertile wild type males. **(b)** Fertilized embryos derived from *brca2 m/m;tp53 m/m* embryos are often morphologically abnormal and arrest at or before sphere stage (approximately four hours post-fertilization). **(c)** At one day post-fertilization, greater than 90% of embryos derived from *tp53 m/m* females are alive and morphologically normal, while all embryos derived from *brca2 m/m;tp53 m/m* females are dead. **(d)** Oocytes from *brca2 m/m;tp53 m/m* females exhibit nuclear abnormalities predominated by aggregated nucleolar material around nuclear margins (black arrowheads). Black box indicates oogonium shown at higher magnification in panel f. **(e)** Oocytes with massive nucleolar condensation are often degenerate. **(f)** Infrequent binucleation occurred in oogonia and oocytes from *brca2 m/m;tp53 m/m* females. Og, oogonia; IA, stage IA oocyte; IB, stage IB oocyte; II, stage II oocyte; III, stage III oocyte. Scale bar = 20 μm.

A total of 3,251 eggs from *tp53 m/m* females and 1,670 eggs from *brca2 m/m;tp53 m/m* females produced from two separate outcrosses were evaluated. Both outcrosses generated a relatively large number of unfertilized eggs (*tp53 m/m* females, n=2,126 eggs; *brca2 m/m;tp53 m/m* females, n=748 eggs). However, *brca2 m/m;tp53 m/m* females generated significantly more inviable eggs (n=822 versus n=180) and significantly fewer fertilized eggs (n=100 versus n=945) compared to *tp53 m/m* females (p<0.0001; **Fig. 1a** and **Table S1**). Embryos derived from *brca2 m/m;tp53 m/m* female zebrafish exhibited a variety of phenotypes during early embryogenesis (**Fig. 1b**). These ranged from apparently normal embryonic cell mass (n=29), reduced cell mass (n=27), abnormally formed cell mass (n=12), or degenerating cell mass (n=32). Regardless of phenotype, all embryos derived from *brca2 m/m;tp53 m/m* zebrafish exhibited an arrest in developmental progression at or before approximately sphere stage (four hours post-fertilization). In comparison, all embryos derived from *tp53 m/m* females exhibited a normal phenotype and did not undergo developmental arrest (n=945). At one day post-fertilization, over 90% (n=874) of embryos derived from *tp53 m/m* females were alive and morphologically normal (**Fig. 1c**). However, no embryos derived *brca2 m/m;tp53 m/m* females survived to one day post-fertilization. These outcomes at one day post-fertilization were statistically significantly different (p<0.0001; **Table S1)**.

Given that BRCA2 is essential for meiotic progression in vertebrate germ cells^25,42,43^, abnormalities in oocyte development might contribute to the phenotype observed in embryos derived from *brca2 m/m;tp53 m/m* zebrafish. We therefore analyzed ovaries from wild type, *tp53 m/m*, and *brca2 m/m;tp53 m/m* zebrafish by histology (n=4 per genotype). Oogonia and oocyte stages were identified as previously described^44^. Ovaries from wild type and *tp53 m/m* zebrafish were histologically similar, and oogonia and oocytes exhibited normal morphology at all stages (**Fig. 1d**). In comparison, meiotic oocytes from *brca2 m/m;tp53 m/m* zebrafish exhibited nuclear abnormalities that were first detectable by histology at stage I and persisted throughout subsequent stages (**Fig. 1d**). Nucleoli were increased in size and decreased in number and were irregularly dispersed around nuclear margins, suggesting aberrant aggregation and distribution of chromosomal material in *brca2 m/m;tp53 m/m* zebrafish oocytes (**Fig. 1d**, arrowheads). Less commonly, *brca2 m/m;tp53 m/m* oocytes exhibited massive consolidation of nuclear material, often in association with oocyte degeneration (**Fig. 1e**). Infrequent binucleation was identifiable in mitotic oogonia and all stages of meiotic oocytes from *brca2 m/m;tp53 m/m* zebrafish (**Fig. 1f**). The extent of nuclear abnormalities in *brca2 m/m;tp53m/m* ovaries was most variable in stage III oocytes (compare panels in **Fig. 1d-f**).

### *brca2* mutation is associated with exhibit aberrant adult cell proliferation in a cancer-prone tissue

To determine *brca2*-associated effects on adult cell phenotypes prior to cancer onset, we focused on the optic nerve pathway (ONP). The natural, well-defined anatomic boundaries of this tissue are ideal for achieving consistent tissue collection, orientation, and histologic sectioning between specimens. Furthermore, the ONP is a predilection site for sarcoma development in both *tp53 m/m* and *brca2 m/m;tp53 m/m* zebrafish^25–27^, with *brca2 m/m;tp53 m/m* zebrafish at significantly increased risk for ocular tumors compared to *tp53 m/m* zebrafish^27^.

We performed a time-course analysis of the ONP in wild type, *tp53 m/m*, and *brca2 m/m;tp53 m/m* between the ages of three and seven months (the mean age at tumor onset in *brca2 m/m;tp53 m/m* zebrafish is 8.7 months^25^). During this period, we noted an abnormality of the choroid rete (CR) in *brca2 m/m;tp53 m/m* zebrafish. The CR is a vascular plexus located subjacent to the retinal choroid that forms a countercurrent capillary system^45^ (**Fig. 2a**). It is derived from the ophthalmic artery and vein and contributes to maintaining oxygen pressure in the retina^46^. In routine hematoxylin and eosin sections, vascular channels of the normal CR are filled with red blood cells and the cellular meshwork forming this structure is largely obscured (**Fig. 2b**). However, the CR in *brca2 m/m;tp53 m/m* zebrafish frequently contained a population of robust spindle cells that were readily apparent between vascular channels (**Fig. 2c**). The incidence of this lesion progressively increased over time (**Fig. 2d**). Notably, this lesion was detectable in all *brca2 m/m;tp53 m/m* specimens after 5.1 months of age. In comparison, incidence was lower and age at onset was higher in *tp53 m/m* zebrafish (**Fig. 2d**). No atypical spindle cells were detected in the CR of wild type zebrafish at any time point examined.

**Figure 2.**
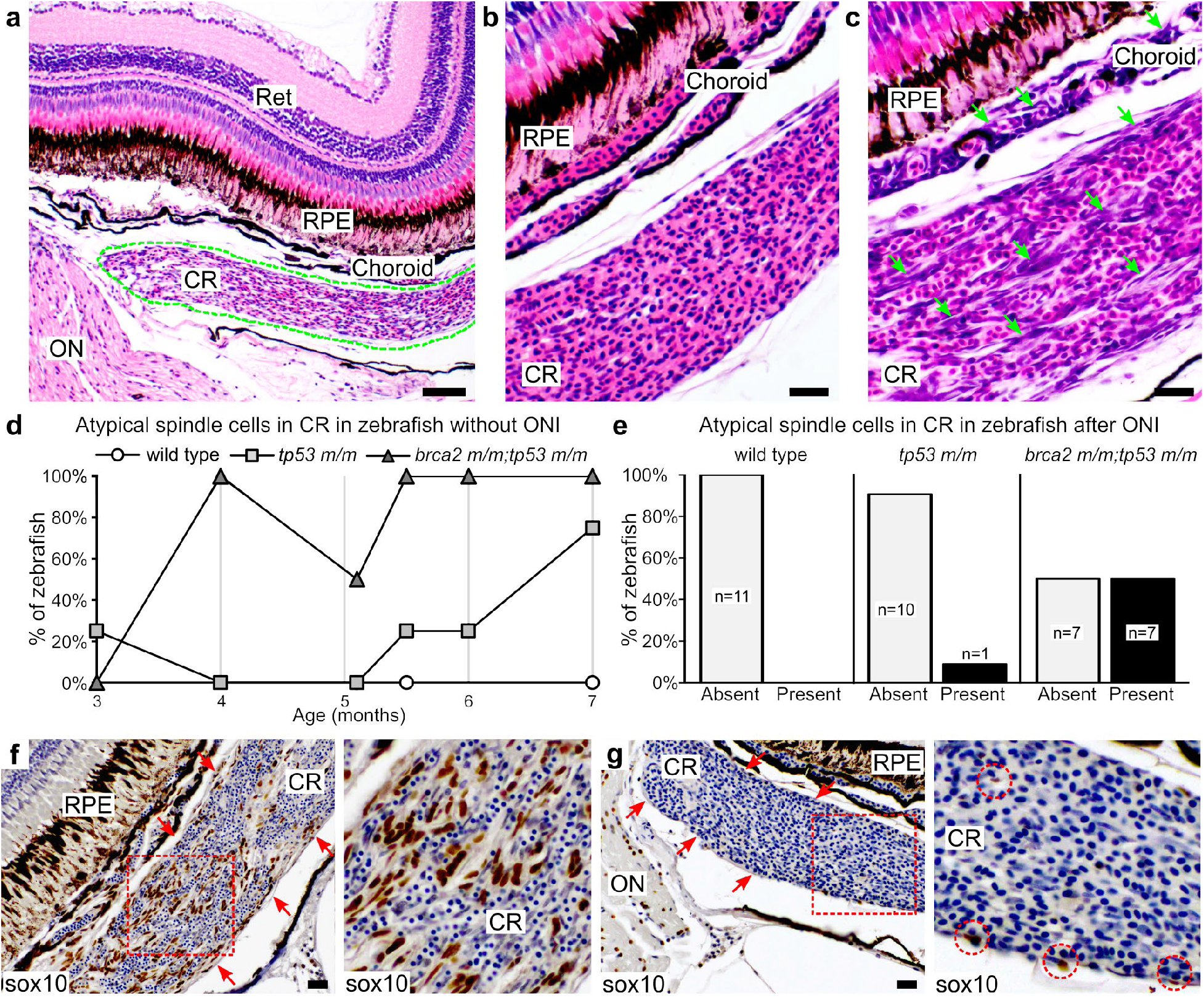
Atypical spindle cells accumulate in the choroid rete of *brca2 m/m;tp53 m/m* zebrafish. **(a)** The choroid rete (outlined in green) is a vascular plexus located subjacent to the choroid. **(b)** Normal choroid rete. **(c)** Choriod rete containing numerous atypical spindle cells (green arrows). **(d)** Numbers of zebrafish that developed atypical spindle cells in the choroid rete over time (n=4 per genotype at each time point analyzed). **(e)** Numbers of zebrafish that received optic nerve injury and developed atypical spindle cells in the choroid rete. **(f)** Atypical spindle cells in the choroid rete are sox10-positive (brown chromogen). Red arrows delineate margins of choroid rete. Area boxed in red is shown at higher magnification to the right. **(g)** Small numbers of sox10-positive cells (brown chromogen) are present in the normal choroid rete. Red arrows delineate margins of choroid rete. Area boxed in red is shown at higher magnification to the right. Red circles identify sox10-positive cells. Ret, retina; RPE, retinal pigmented epithelium; CR, choroid rete; ON, optic nerve. Scale bar = 50 μm (panel a); 20 μm (panels b, c, f, h).

### Optic nerve injury (ONI) induces an enhanced proliferative response in *brca2 m/m;tp53 m/m* zebrafish

As described above, the ONP is a cancer predilection site in *tp53 m/m* and *brca2 m/m;tp53 m/m* zebrafish^25–27^ and is also notable for unique properties that support complete regeneration of the injured retina and optic nerve^32,33,36–38^. We therefore sought to determine how proliferative and neoplastic phenotypes in the ONP might relate to injury response and regenerative capacity. We first assessed the short-term effects of ONI by performing unilateral ONI in wild type (n=11), *tp53 m/m* (n=11), and *brca2 m/m;tp53 m/m* (n=14) zebrafish and assessing the injury response at three days and two weeks post-injury (**Fig. 3a-c, Tables S2-S3**). At both three day and two week time points, the total cellularity of the injured optic nerve was significantly increased compared to the uninjured optic nerve in *brca2 m/m;tp53 m/m* zebrafish (p=0.0001 and p<0.0001, respectively; **Fig. 3d,e** and **Table S2**). In comparison, total cellularity was significantly increased in wild type and *tp53 m/m* zebrafish only at two weeks post-injury (p=0.0464 and p=0.0130, respectively; **Fig. 3d,e** and **Table S2**). Comparison between genotypes indicated that the injury effect in *brca2 m/m;tp53 m/m* was significantly greater than in wild type or *tp53 m/m* cohorts at both three days and two weeks post-injury (**Fig. 3d,e** and **Table S2**).

**Figure 3.**
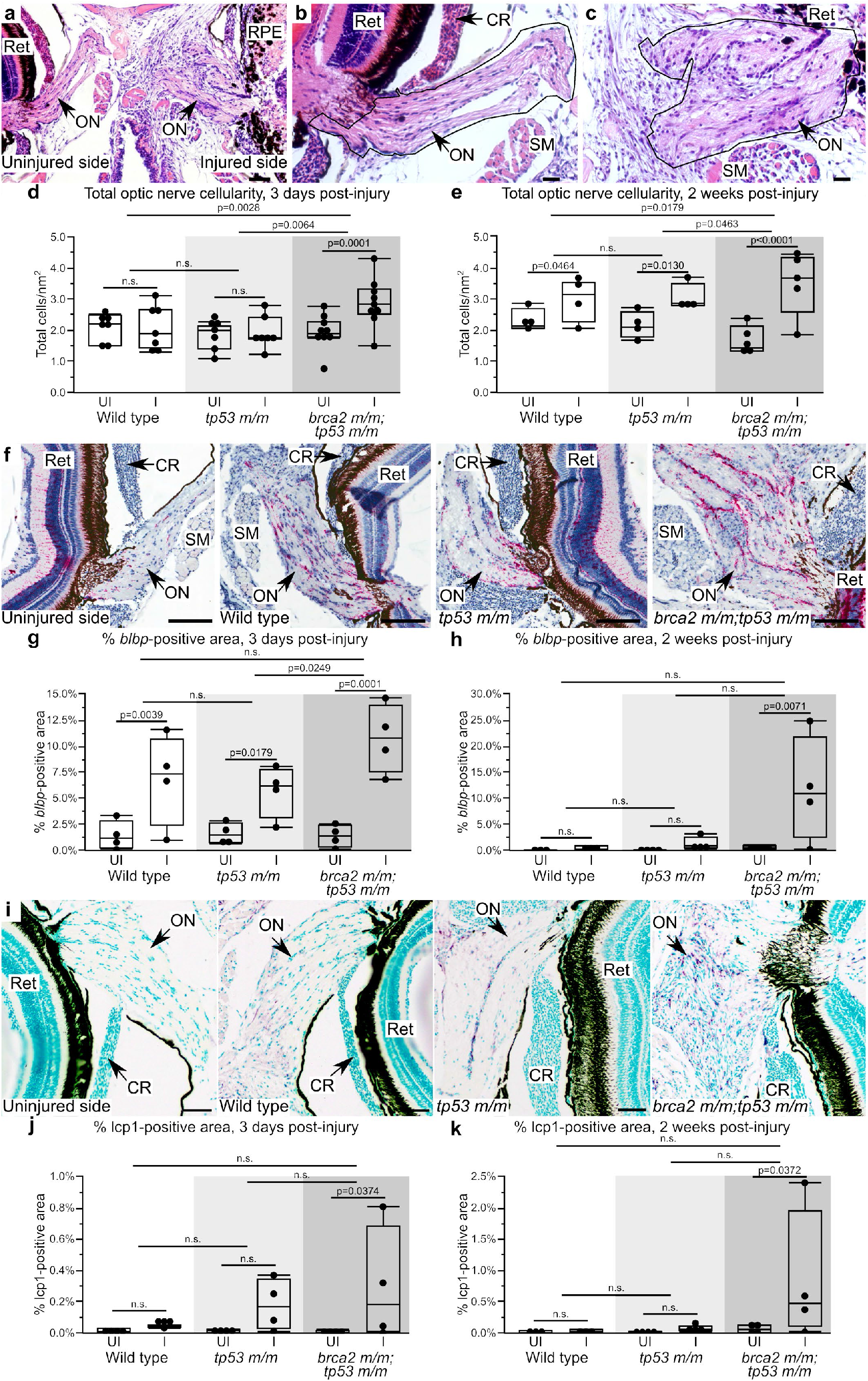
*brca2 m/m;tp53 m/m* zebrafish exhibit increased proliferative responses in the injured optic nerve. **(a)** Representative histologic section of the optic nerve pathway after unilateral optic nerve injury. **(b,c)** Representative example of the uninjured (b) and injured (c) optic nerve. The portion of the nerve used for quantification of total cellularity is outlined in black. **(d,e)** Quantitative analysis of the total cellularity in the uninjured versus injured optic nerve three days (d) and two weeks (e) post-injury. **(f)** Representative examples of *blbp* expression (red chromogen), a marker for radial glial cells, in the uninjured and injured optic nerves. **(g,h)** Quantitative analysis of *blbp*-positive area in the uninjured versus injured optic nerve three days (g) and two weeks (h) post-injury. **(i)** Representative examples of lcp1 expression (purple chromogen), a marker for monocytes/macrophages, in the uninjured and injured optic nerves. **(j,k)** Quantitative analysis of lcp1-positive area in the uninjured versus injured optic nerve three days (j) and two weeks (k) post-injury. Ret, retina; SM, skeletal muscle; CR, choroid rete; ON, optic nerve; UI, uninjured; I, injured. Scale bar = 50 μm (panels a, i); 20 μm (panels b, c); 100 μm (panel f).

To identify specific cell types that contribute to the increased cellularity observed in injured optic nerves, we performed a series of quantitative analyses by immunohistochemistry and *in situ* hybridization (details of statistical analyses are in **Tables S2** and **S3**). First, we analyzed injured and uninjured optic nerves from four zebrafish at each time point (three days and two weeks post-injury) for the presence of stem and progenitor cell populations. We assessed the expression of *blbp*, a marker for radial glial cells^47^ **(Fig. 3f)**; sox2, a marker for neural stem cells^48^ **(Fig S2a)**; and sox10, a marker for neural crest progenitor cells^49,50^, oligodendrocytes and oligodendrocyte precursors^51^, and Schwann cells and Schwann cell precursors^52^ **(Fig S2d)**. As *blbp* expression is cytoplasmic, we could not reliably identify and count individual *blbp*-expressing cells. We therefore quantified the total area of *blbp* expression in the optic nerve. At three days post-injury, *blbp* expression was significantly increased in the injured optic nerve compared to the uninjured optic nerve in all cohorts (wild type, p=0.0039; *tp53 m/m*, p=0.0179; *brca2 m/m;tp53 m/m*, p=0.0001; **Fig. 3g**). *brca2 m/m;tp53 m/m* exhibited a sustained and significant increase in *blbp* expression at two weeks post-injury that was not observed in wild type or *tp53 m/m* zebrafish (p=0.0071; **Fig. 3h**). On the other hand, neither sox2-expressing cells (**Fig. S2b,c**) nor sox10-expressing cells (**Fig. S2e,f**) appeared to contribute to the significant increases in cellularity observed in the injured optic nerve in *brca2 m/m;tp53 m/m* zebrafish. There were generally no significant differences observed in comparisons between genotypes, indicating that the significant injury effect observed in *brca2 m/m;tp53 m/m* zebrafish based on total cellularity (**Fig. 3d, e**) was not attributable to a single cell type.

Next, we analyzed injured and uninjured optic nerves for the presence of inflammatory and reactive cell populations. We assessed the expression of lcp1, a marker for monocytes and macrophages^53^ (**Fig. 3i**), and krt18, a marker for reactive astrocytes in the ONP after injury^54^ (**Fig S2g**). ln addition to monocyte/macrophage populations, lcp1 is reportedly expressed by microglial cells in zebrafish^55^. However, we were unable to detect lcp1-positive microglial cells in sections of zebrafish brain and therefore considered lcp1 as a marker for monocytes and macrophages. Both lcp1 and krt18 are expressed in the cytoplasm, and therefore the total area of lcp1 or krt18 expression was quantified similar to *blbp* expression. At both three days and two weeks post-injury, lcp1 expression was significantly increased only in the injured optic nerve in *brca2 m/m;tp53 m/m* zebrafish (p=0.0374 and p=0.0372, respectively; **Fig. 3j,k**). In comparison, krt18 expression were not significantly different in the injured versus uninjured optic nerves at most time points for any of the three cohorts (**Fig. S2h,i**).

Finally, we assessed the CR in zebrafish that received ONI for the presence of aberrant spindle cells as observed in uninjured zebrafish. This cell population was present in 7 of 14 *brca2 m/m;tp53 m/m* zebrafish versus 1 of 11 *tp53m/m* zebrafish, and was not identified in any wild type zebrafish (**Fig 2e**). When present, atypical spindle cells were identified at both three days and two weeks post-injury and there was no clear predilection for the injured or uninjured side. We further found that the atypical spindle cells identified in the CR were uniformly sox10-positive and were distributed both on the periphery and within the body of the CR (**Fig. 2f**). These expression patterns were similar regardless of whether the spindle cell population had arisen on the injured or uninjured side. In the normal CR, low numbers of sox10-positive cells were present in the CR and were largely confined to the periphery (**Fig. 2g**).

### ONI does not significantly increase ocular tumorigenesis, but affects the incidence and sidedness of ocular lesions in *brca2 m/m;tp53 m/m* versus *tp53 m/m* zebrafish

To determine how injury and regenerative responses affect ocular tumorigenesis in zebrafish, we analyzed tumor development in *brca2 m/m;tp53 m/m* and *tp53 m/m* zebrafish that received unilateral ONI at three months of age. The ONI group was compared to *brca2 m/m;tp53 m/m* and *tp53 m/m* zebrafish from a previously reported cohort, designated as the control group^27^ (see Methods and **Table S4** for details). Tumor development in ONI and control groups is summarized in **Table 1**. First, we compared the proportion of ocular versus non-ocular tumors in the ONI group versus the control group and in ONI and control groups segregated by genotype. In each comparison, ONI was not associated with an increased proportion of ocular tumors (**Table 1** and **Table S1**). Next, we compared the side of ocular tumor development in ONI versus control groups to determine whether there was a side predilection for ocular tumorigenesis in either population. In each comparison, there was no significant difference in the proportions of ocular tumors arising on the right side (ONI side), left side (non-ONI side), or bilaterally in ONI versus control groups (**Table 1** and **Table S1**).

**Table 1.** Characteristics of tumor development in zebrafish receiving optic nerve injury (ONI) versus control zebrafish.

| **Location of tumor development (ocular versus non-ocular)** |  |  |  |
| --- | --- | --- | --- |
|  | **Ocular tumors** | **Non-ocular tumors** | **Total tumors** |
| **ONI group** | **45 (37%)** | **77 (63%)** | **122** |
| *brca2 m/m;tp53 m/m* | 29 (43%) | 39 (57%) | 68 |
| *tp53 m/m* | 16 (30%) | 38 (70%) | 54 |
| **Control group** | **36 (34%)** | **70 (66%)** | **106** |
| *brca2 m/m;tp53 m/m* | 23 (42%) | 32 (58%) | 55 |
| *tp53 m/m* | 13 (25.5%) | 38 (74.5%) | 51 |
| **Side of ocular tumor development** |  |  |  |
|  | **Right/ONI side** | **Left /non-ONI side** | **Bilateral** |
| **ONI group** | **18 (40%)** | **19 (42%)** | **8 (18%)** |
| *brca2 m/m;tp53 m/m* | 13 (45%) | 10 (34%) | 6 (21%) |
| *tp53 m/m* | 5 (31%) | 9 (56%) | 2 (13%) |
| **Control group** | **17 (47%)** | **14 (39%)** | **5 (14%)** |
| *brca2 m/m;tp53 m/m* | 11 (48%) | 9 (39%) | 3 (13%) |
| *tp53 m/m* | 6 (46%) | 5 (39%) | 2 (15%) |

We have previously shown that both ocular and non-ocular tumors from *brca2 m/m;tp53 m/m* and *tp53 m/m* zebrafish are predominantly sarcomas that exhibit histologic and immunohistochemical features consistent with malignant peripheral nerve sheath tumor^25,29^. The immunohistochemical expression profile of these tumors is not affected by *brca2* genotype^29^. Ocular tumors arising in both the ONI and control cohorts were histologically similar and consistent with our previous identification of these tumors as sarcomas with features of malignant peripheral nerve sheath tumor. To further characterize these tumors, we analyzed a subset of ocular tumors from ONI and control zebrafish for expression of *blbp*, sox2, and sox10 (**Fig. 4a** and **Fig. S3**). Expression of these markers was similar in tumors derived from ONI and control cohorts. Semi-quantitative analysis of marker expression demonstrated that most tumors exhibited little or no expression of either *blbp* or sox2 (**Fig S3**). However, tumors from both ONI and control groups exhibited strong and ubiquitous nuclear sox10 expression (**Fig. 4a**).

**Figure 4.**
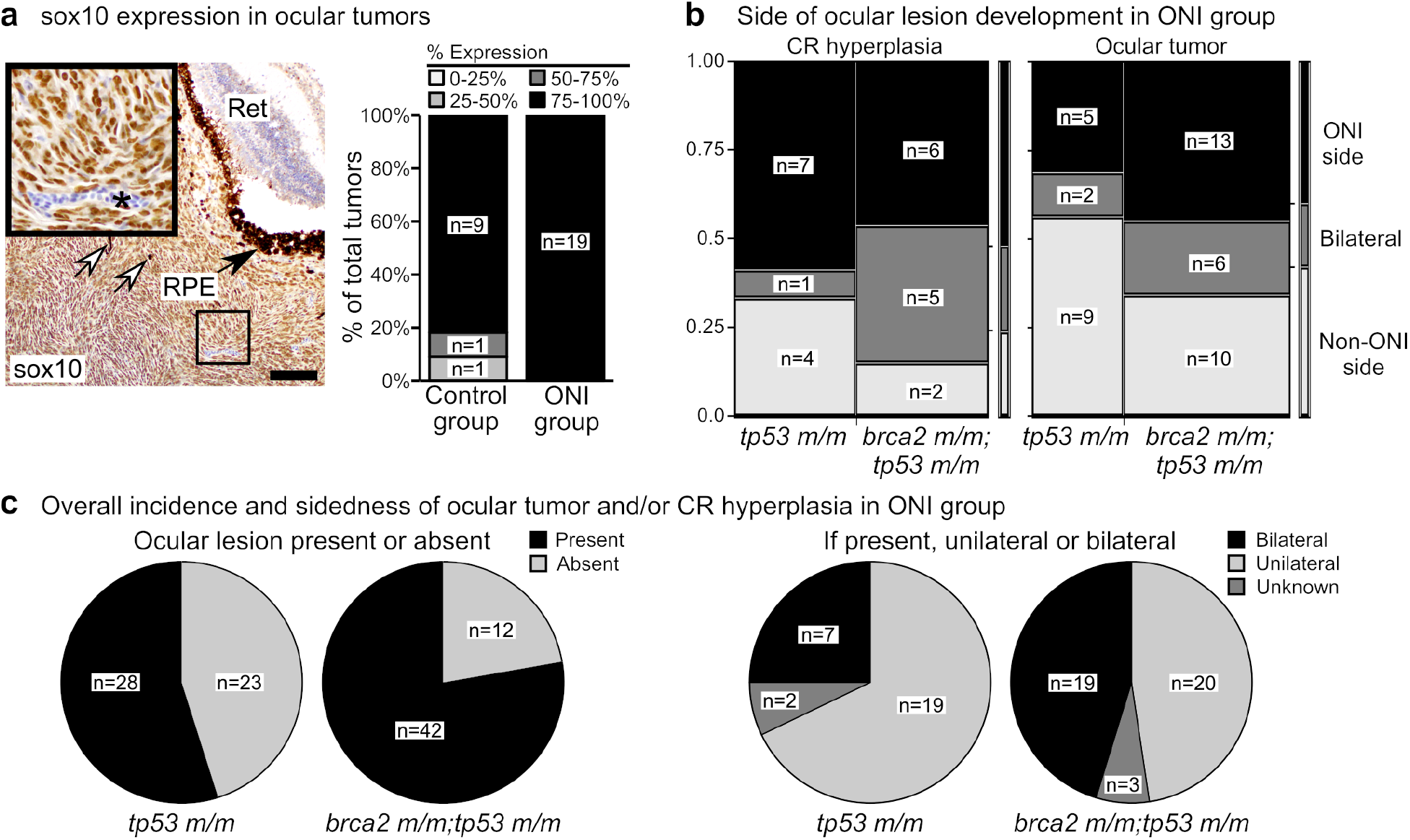
Optic nerve injury (ONI) and *brca2* genotype exert variable effects on the development of proliferative and neoplastic lesions in the optic nerve pathway. **(a)** The majority of zebrafish ocular tumors highly express sox10 regardless of ONI status. White arrows indicate dispersed melanin pigment within tumors. Inset shows tumor cells with positive nuclear sox10 expression (brown chromogen). Asterisk indicates a blood vessel containing nucleated erythrocytes that do not express sox10. **(b)** In zebrafish that received ONI, most *brca2 m/m;tp53 m/m* zebrafish developed atypical spindle cells in the choroid rete (CR) on the injured side (ONI side) or bilaterally, in contrast to ocular tumor development. **(c)** In zebrafish that received ONI, *brca2 m/m;tp53m/m* zebrafish were more likely to develop ocular lesions (ocular tumor and/or hyperplastic spindle cells in the choroid rete) and these lesions were more likely to be bilateral. HE, hematoxylin and eosin; Ret, retina; RPE, retinal pigmented epithelium; ONI, optic nerve injury; CR, choroid rete. Scale bar = 50 μm.

Since we had determined that *brca2 m/m;tp53 m/m* frequently develop an atypical spindle cell population in the choroid rete of the eye between 3 and 7 months of age (**Fig. 2**), we investigated the incidence of this lesion in older animals that were followed for tumor development. Because cross-sections of the head were not routinely collected from the control group (derived from a previous study not specifically focused on the ONP; see Methods), our analysis was limited to zebrafish from the ONI group. We first assessed the incidence of atypical spindle cells in the CR in zebrafish from the ONI group that did not develop ocular tumors. Similar to earlier time points, we found that a higher proportion of *brca2 m/m;tp53 m/m* zebrafish (n=13 of 25, 52%) exhibited CR atypical spindle cells compared to *tp53 m/m* zebrafish (n=12 of 35, 34%), although this increase was not statistically significant (**Table S1**).

We next compared the side for development of ocular tumors or CR atypical spindle cells in *brca2 m/m;tp53 m/m* zebrafish versus *tp53 m/m* from the ONI group (**Fig. 4b**). There was no significant difference in the proportions of ocular tumors or CR atypical spindle cells arising on the ONI side, non-ONI side, or bilaterally between genotypic groups (**Table S1**). However, we noted two differences in ocular tumorigenesis versus CR atypical spindle cells in these analyses. First, CR atypical spindle cells in both *tp53 m/m* and *brca2 m/m;tp53 m/m* zebrafish were more frequently identified on the ONI side compared to the non-ONI side, in contrast to ocular tumors (**Fig. 4b**). Second, *brca2 m/m;tp53 m/m* zebrafish exhibited a relatively greater proportion of bilateral CR atypical spindle cells compared to the proportion of bilateral ocular tumors (**Fig. 4b**). Lastly, we compared the overall incidence and sidedness of ocular lesions (ocular tumor or CR atypical spindle cells) in *brca2 m/m;tp53 m/m* zebrafish versus *tp53 m/m* from the ONI group (**Fig. 4c**). After ONI, *brca2 m/m;tp53 m/m* zebrafish were significantly more likely to exhibit an ocular lesion than *tp53 m/m* zebrafish, and ocular lesions, when present, were more often bilateral in *brca2 m/m;tp53 m/m* zebrafish (p=0.0220 and p=0.1207, respectively; **Table S1**).

## Discussion

The exploration of noncancerous cell phenotypes associated with mutations in cancer susceptibility genes can provide important insights into how such mutations affect cell behaviors and responses to stimuli. In the current study, we analyzed several noncancerous cell phenotypes linked to BRCA2 mutation using a zebrafish model. BRCA2 is required for error-free resolution of double-strand DNA breaks by homologous recombination in both mitotic and meiotic cells, and also participates in processes such as replication fork protection and R loop processing (reviewed in ^56–58^). Heritable BRCA2 mutations are associated with significantly increased risk for several cancer types in humans, including breast, ovarian, prostate, and pancreatic cancer^56^, and impaired capacity for homologous recombination has been identified more broadly across multiple human cancers (“BRCAness”)^59–61^.

The processes of embryonic development and the inflammatory/injury response can be partially recapitulated during carcinogenesis, as molecular and cellular programs that are activated during these processes can be co-opted by cancer cells^1–5^. Furthermore, these normal processes can be influenced by cancer-associated genetic mutations and thereby directly contribute to cancer initiation and progression. For example, pancreatic cancer can be induced by the cooperating effects of *KRAS* mutation, inflammation, and tissue injury^7^, which has been attributed to specific epigenetic alterations that are uniquely driven by these combined genetic and microenvironmetal factors^62^. Cancer initiation is also associated with the activation of molecular signaling pathways that normally function during embryogenesis. However, developmental effects caused by mutation or loss of cancer-associated genes are often very different than adult phenotypes, as exemplified by early embryonic lethality versus adult cancer susceptibility^11–22^.

Determining how cancer-causing genetic mutations affect adult versus embryonic cell populations is likely to reveal distinct signaling pathways that underlie these phenotypes and are of significant relevance to cancer initiation (i.e., cell proliferation versus cell death). Unfortunately, early embryonic lethality in mouse models for cancer-associated genes such as BRCA2 is a confounding factor. In these cases, zebrafish can provide an excellent complementary model, as they exhibit conserved genetic susceptibility to cancers for many genes^63–68^. The *brca2*-mutant zebrafish model is fully viable in the homozygous condition and captures the collaborative effects of *brca2* and *tp53* mutations in carcinogenesis that characterize human BRCA2-associated cancers^25,27,28^. We thus used this model system to determine how *brca2* mutation affects cell phenotypes during embryogenesis and in the response to tissue injury. As the optic nerve pathway (ONP) is a predilection site for cancer development *in brca2 m/m;tp53 m/m* zebrafish, evaluations of the injury response in adult animals focused on this tissue.

We first assessed the role for maternally provided mRNA for *brca2* during early embryonic development in zebrafish. The maternal-to-zygotic transition is characterized by the degradation of maternal mRNA and onset of zygotic gene activation (ZGA)^69^. While mice initiate ZGA at the 1-cell stage and clear most maternal mRNA by the 2-cell stage, zebrafish do not undergo these processes until the mid-blastula transition at cleavage cycle 10^69–71^. These differences in availability of maternally provided transcripts may be a factor in survival of *brca2*-mutant zebrafish embryos. We have previously shown that zebrafish embryos possess abundant maternal mRNA for *brca2^25^*, and *brca2* is both maternally and zygotically expressed in early-stage embryos^70^. In accordance with this, we show here that zebrafish embryos lacking maternally provided mRNA for full-length brca2 exhibit developmental arrest and death at approximately mid-blastula stage. However, detailed analysis of ovaries from female *brca2 m/m;tp53 m/m* zebrafish reveal abnormalities in developing oocytes that could also contribute to this embryonic phenotype. These include both aberrant localization of nuclear content and evidence for cytokinetic defects; the former observation has been reported in another *brca2*-mutant zebrafish model^72^. Beyond the canonical role for BRCA2 in dsDNA break repair, *in vitro* analyses in mitotic cells indicate that BRCA2 participates in cytokinesis^73^ and chromosomal alignment/segregation^74,75^. Less is known about the role for BRCA2 in meiosis due to the difficulty in establishing *Brca2*-knockout mouse models, although meiosis-specific binding partners required for BRCA2 localization to chromosomes were recently characterized^76–78^. However, mouse models with oocyte-specific reduction in Brca2 expression^43^ or Brca2 deletion^42^ displayed nuclear abnormalities suggesting errors in chromosomal localization in oocytes. Loss of functional BRCA2 is likely to disrupt meiotic progression and early embryonic development through multiple mechanisms. Further studies will be required to segregate and clarify the functions of BRCA2 in meiotic oocytes versus early-stage embryos.

We next determined that loss of functional brca2 in adult zebrafish induces aberrant proliferative responses in the ONP, which is a highly cancer-prone tissue in *brca2 m/m;tp53 m/m* zebrafish. We identified an anomalous spindle cell population that was highly prevalent in the choroid rete of *brca2 m/m;tp53 m/m* zebrafish in the unperturbed ONP prior to cancer onset. Uniform sox10 positivity suggests that the choroid rete spindle cells are of neural crest origin, oligodendroglial, or Schwann cell origin. Our current and prior^29^ immunohistochemical analyses of ocular tumors in *brca2 m/m;tp53 m/m* zebrafish are consistent with malignant peripheral nerve sheath tumor (MPNST) and demonstrate widespread sox10 expression in tumors, supportive of Schwann cell origin. The zebrafish choroid, and presumably the choroid rete, contains small myelinated nerve processes that are the likely source for Schwann cells in this location^79^. We therefore hypothesize that tumors in the optic nerve pathway in our model arise from this aberrantly proliferative Schwann cell population. In comparison, conditional deletion of *Brca2* in mouse prostatic epithelium induces epithelial hyperplasia and low-grade prostate intraepithelial neoplasia (PIN) that is exacerbated by concurrent *Tp53* mutation^80^. On the other hand, *Brca2* knockout in mouse T lymphocytes causes a decline in T cell numbers over time^81^. These data from zebrafish and mouse models suggests that BRCA2 mutation or loss affects different cell types differently, and can enhance the growth of certain noncancerous cell populations in specific tissues/contexts. An important next step will be to determine why a particular microenvironment promotes cell proliferation and subsequent cancer initiation in the context of heritable *BRCA2* mutation.

We subsequently assessed the injury response in the cancer-prone ONP, since numerous studies have demonstrated key similarities between injury responses and cancer progression at the molecular, cellular, and tissue level^1–3,6^. The ONP in zebrafish is uniquely supportive of complete optic nerve regeneration due in part to a permissive microenvironment that supports axonal regrowth. We therefore speculated that cellular responses to optic nerve injury (ONI) might differ in cancer-prone versus non-cancer-prone individuals, and that cancer predisposition might be related to regenerative capacity in this pro-growth environment. In short-term studies, the proliferative response to ONI was significantly enhanced in *brca2 m/m;tp53 m/m* zebrafish compared to *tp53 m/m* or wild type cohorts. This included both progenitor cells (radial glia) and inflammatory cells (monocytes/macrophages); other cell populations that were not investigated here may have added to the overall increase in cellularity. Interestingly, cardiomyocyte proliferation during heart regeneration in zebrafish increases in the context of homozygous *tp53* mutation^82^. Similar p53-associated effects on cell proliferation are described during early stages of limb regeneration in salamanders^83^. We are not aware of any studies that test the role for BRCA2 in vertebrate regeneration; however, the orthologue for BRCA2 contributes to axonal regeneration in the nematode *Caenorhabditis elegans*^84^. We also noted some differences in cellular responses to ONI in the current study, e.g., sox10-expressing cells, compared to other reports of optic nerve injury in fish^85,86^. These differences reflect variations in which portion of the injured optic nerve was analyzed, and may also be affected by differences in analytical time points. Together these studies indicate that injury responses are significantly altered by mutations in cancer-associated genes in vertebrate animals.

The potential for cancers to arise from regenerating cell populations in vertebrates is variable. In salamanders and newts, regenerative tissues are highly resistant to chemical carcinogenesis and malignant transformation is suppressed^87–89^. Here we found no direct effect (positive or negative) on tumorigenesis following ONI in zebrafish with heritable *brca2* and *tp53* mutations. However, we did note that bilateral ocular lesions were more common in *brca2 m/m;tp53 m/m* zebrafish after ONI than might be expected based on the incidence of bilateral ocular tumors in controls. In contrast, a zebrafish model for *KRAS^G12V^*-driven melanoma subjected to repeated cycles of tail amputation and regeneration developed melanoma at the resection site^90^. The differences in regeneration-associated tumorigenesis in *KRAS^G12V^* zebrafish versus *brca2*-mutant/*tp53*-mutant zebrafish could reflect the relative impact of chronic repeated injuries versus a single injury event on tumor initiation. Alternatively, differences in the regenerative process might affect tumorigenic potential in zebrafish. While ONI is resolved by regeneration of axonal fibers from surviving retinal ganglion, tail resection is resolved by the more complex process of epimorphic regeneration^31^. Epimorphic regeneration requires repatterning and regrowth of multiple tissue types and is achieved via dedifferentiation and subsequent redifferentiation of mature cell populations^91^.

In summary, we demonstrate that phenotypes linked to BRCA2 mutation in mammals, ranging from early embryonic death to cancer predisposition, are captured in the *brca2*-mutant zebrafish model. We find that *brca2*-associated embryonic lethality is likely to reflect a combination of cellular defects that arise during both mitosis (oogonia, embryos) and meiosis (oocytes). We also identify expansion of several adult cell populations arising under basal conditions and during the post-injury response in a tissue at high risk for cancer onset, including a putative precancerous population. These studies confirm stage- and context-dependent roles for BRCA2 in cell survival and growth that are highly relevant to BRCA2-associated carcinogenesis.

## Materials and Methods

### Zebrafish study cohorts

Experiments were performed with adult wild type (AB) zebrafish and adult zebrafish from the *brca2^hg5^* and *tp53^zdf1^* mutant zebrafish lines, corresponding to *brca2^Q658X^* and *tp53^M214K^* mutations^25,26^. Mutant alleles are hereafter referred to as “m”. Details of the experimental groups are included in **Table S4**. Groups including *tp53 m/m* and *brca2 m/m;tp53 m/m* were comprised of siblings genotyped for presence or absence of the *brca2^Q658X^* mutation. For analysis of oocyte morphology, groups consisted of age-matched wild type, *tp53 m/m*, and *brca2 m/m;tp53 m/m* female zebrafish. For analyses of injury response and tumorigenesis, the ONI group consisted of age-matched cohorts, and zebrafish with *brca2* and *tp53* mutations were siblings derived from two clutches. For analysis of tumorigenesis, the control group consisted of zebrafish siblings with *brca2* and *tp53* mutations derived from a single clutch. The control group was previously described in a separate study analyzing tumor ploidy^27^. All animal studies were approved by the Institutional Animal Care and Use Committee, North Carolina State University, Raleigh, NC, performed in accordance with approved protocols, and complied with ARRIVE guidelines.

### Zebrafish husbandry and genotyping

Zebrafish used in this study were raised as previously described27 on a Pentair Z-Hab Duo recirculating aquaculture system and maintained on a 14-hour light/10-hour dark cycle. The zebrafish colony undergoes routine sentinel testing for infectious organisms and is negative for known zebrafish pathogens. Zebrafish were monitored for gross evidence of tumor development and humanely euthanized with Tricaine methanesulfonate (300 mg/L) in system water buffered with Sodium Bicarbonate to a pH of ~ 7.0 when tumors were visibly apparent. Live adult zebrafish were genotyped for the *brca2^Q658X^* mutation at three months of age by sequencing over the mutation site as previously described^28^. Zebrafish with *tp53^zdf1^* mutation were maintained as a homozygous mutant line.

### Optic nerve injury

Optic nerve injury was performed on randomly selected and anesthetized 5-month-old zebrafish placed in left lateral recumbency on a wet sponge under a stereomicroscope. The right eye was gently displaced, and the optic nerve was crushed with micro-forceps. All zebrafish received equivalent injury and were recovered in system water. Zebrafish were collected and euthanized at 3 days post-injury for short-term analyses and at 2 weeks after injury for long-term analyses. Zebrafish were observed for tumor development for up to 19 months of age.

### Tissue collection, immunohistochemistry, and *in situ* hybridization

Zebrafish from the ONI group were decapitated caudal to the gills after humane euthanasia was performed as described above. Heads were placed in 4% paraformaldehyde for 18-24 hours, decalcified in 12% EDTA for two days, and transferred to 70% ethanol. If zebrafish were to be used for tumorigenesis studies, the body was similarly processed. Tissue specimens were embedded in paraffin, and unstained or hematoxylin and eosin stained transverse sections were prepared by the Histology Laboratory, NC State University, College of Veterinary Medicine so that both optic nerves were in the plane of section. Zebrafish tissues from the tumor-bearing control group and from female wild type, *tp53 m/m*, and *brca2 m/m;tp53 m/m* zebrafish were collected and processed as previously described^27^.

Immunohistochemistry on paraffin-embedded sections was performed for expression of zebrafish sox2, sox10, lcp1, and krt18 as previously described^29^ with minor modifications. Antibodies used included rabbit anti-SOX2 (Abcam ab97959); rabbit anti-SOX10 (GeneTex GTX128374); rabbit anti-lcp1 (Genetex GTX134697); and mouse anti-KRT18 (Abcepta AT2655a). Detection was achieved with ImmPACT DAB peroxidase substrate (sox2, sox10; Vector #SK-4105) or ImmPACT VIP peroxidase substrate (lcp1, krt18; Vector Labs #SK-4605) and sections were counterstained with Mayer’s hematoxylin (sox2, sox10) or methyl green (lcp1, krt18). ACD RNAscope RNA *in situ* hybridization was performed according to manufacturer specifications to determine expression of zebrafish *blbp* (ACD probe 414651). The RNAscope 2.5 HD Assay - RED (ACD 322350) was used for detection. The brain and retina served as internal positive controls for sox2, sox10, and *blbp* expression (**Fig. S1**). A sample of zebrafish spleen served as a positive control for lcp1 expression (**Fig. S1**). As krt18 is reportedly only expressed by reactive astrocytes in response to injury^54^, there was no additional positive control tissue other than the injured optic nerve in tissue specimens. Negative controls included slides incubated with secondary antibody only and slides incubated with an RNA probe against Bacillus subtilis dihydrodipicolinate reductase (*dapB)* (**Fig. S1**).

### Imaging and image analysis

For quantitative analyses of ONI specimens, slides were scanned at 20X magnification to generate 5976 x 7740 digital images (Translational Pathology Lab, University of North Carolina) or imaged at 20X magnification on an Olympus brightfield microscope with Olympus cellSens Imaging Software. The injured nerve and contralateral uninjured nerve were analyzed for each specimen. Quantitative analyses were performed using either a single digital image or multiple aligned digital images captured from hematoxylin and eosin-stained sections. Images were minimally and globally processed with the GNU Image Manipulation Program, version 2.8.6 (http://www.gimp.org/) and the line tool was used to outline the optic nerve area in each tissue section. Quantitation of total cellularity, total positive cells (sox2, sox10), or percent positive area (*blbp*, lcp1, krt18) was performed with ImageJ, using the outlined area for each optic nerve to define a region of interest. Total cellularity was determined using the ImageJ Fiji Cell Counter tool (https://fiji.sc/) (ref). Red blood cells were excluded from cell counts based on their appearance as ovoid, nucleated cells with brightly eosinophilic cytoplasm. Expression of *blbp*, sox2, sox10, lcp1, and krt18 were determined with the IHC Toolbox plugin for ImageJ (https://imagej.nih.gov/ij/plugins/). A positive control specimen was used during the training process to generate a model that identified positive pixels in digital images for each marker. Since there was no positive control for krt18, the model generated for lcp1 expression was used because krt18 and lcp1 expression were detected with the same chromogen. sox2 and sox10 expression were determined by quantifying the total cells versus sox2-or sox10-expressing cells, based on nuclear expression of these markers, and calculating the ratio of positive cells to total cells within the outlined nerve. *blbp*, lcp1, and krt expression were determined by quantifying the percent positive area within the total area of the outlined nerve.

Histologic analyses and imaging of tissue specimens were performed by a board-certified veterinary pathologist (HRS) using an Olympus BX43 light microscope with Olympus DP27 digital camera and Olympus cellSens Imaging Software. Images were minimally and globally processed with the GNU Image Manipulation Program, version 2.8.6 (http://www.gimp.org/). Semi-quantitative analyses of sox2, sox10, and *blbp* expression were performed in a subset of tumors arising on the right (ONI) side and left (non-ONI) side from optic nerve injury and control cohorts. Tumor expression of these markers was analyzed and scored with an Olympus BX51 light microscope by a single investigator (VAK). Marker expression was scored as a percentage of total tumor tissue in each section (0-25%, 25-50%, 50-75%, or 75-100%) by visual assessment of the entire tumor at 40X, 100X, and 200X magnification.

### Embryo phenotyping

Embryos were derived from *tp53 m/m* and *brca2 m/m;tp53 m/m* female zebrafish outcrossed to fertile wild type males in two independent experiments. Zebrafish were maintained overnight in breeding chambers without dividers in groups of three to four females per two males, and eggs were collected the following morning upon cessation of breeding behavior. Every egg derived from each clutch was assessed using a Nikon SMZ1000 stereomicroscope and counted as either fertilized (intact egg undergoing cell division), unfertilized (intact egg without cell division), or inviable (degenerate egg). Fertilized eggs were sorted into 100 mm Petri dishes in egg water (60 ug/ml “Instant Ocean” sea salts and 0.0002% methylene blue in distilled water) at a density of up to 55 embryos per dish and incubated at 28°C degrees in a dedicated incubator. Embryos were periodically observed at zero days post-fertilization to assess developmental progress. At one day post-fertilization, embryos were scored as exhibiting normal phenotype, abnormal phenotype, or inviable using established staging criteria^92^. For one group of embryos derived from *tp53 m/m* females, 35 fertilized embryos were removed from a total of 522 fertilized embryos on day 0 for an unrelated experiment and are not included in the total on day 1.

### Criteria for exclusion

Individual zebrafish that were (1) found dead; (2) lost the right eye after injury or (3) had histological evidence of unusually severe tissue damage after ONI were excluded from analysis. In addition, specimens for which both optic nerves or both choroid rete could not be identified in tissue sections were excluded from the relevant analyses. See **Table S4** for additional details.

### Statistical Analyses

Statistical analyses were performed using SAS software version 9.4 (SAS Institute Inc., Cary, NC) with statistical significance set at an alpha value of p ≤ 0.05. Comparisons of cellularity and marker expression (sox2, sox10, *blbp*, lcp1, krt18) were performed using a mixed effect model. Fisher’s Exact test was used to compare population proportions for the following assessments: tumor location; sidedness of ocular tumors; presence of atypical spindle cells in the choroid rete; presence of any ocular lesion; sidedness of ocular lesions. Details of statistical analyses and outcomes are shown in **Table S1-S3**.

## Acknowledgements

This work was supported in part by the Office of Research Infrastructure Programs of the National Institutes of Health under award number K01OD021419 and by the NCSU Faculty Research and Professional Development Fund under award number 2015-2934. The content is solely the responsibility of the authors and does not necessarily represent the official views of the National Institutes of Health. The authors would like to thank Ms. Laura Miller (NC State University Histology Laboratory) for preparing unstained and hematoxylin and eosin stained specimens from zebrafish tissues.

## Author Contributions

H.R.S designed the research; V.A.K, A.A.S., J.L.F., X.M., and H.R.S performed research and analyzed data; V.A.K. and H.R.S wrote the manuscript.

## Additional Information

The author(s) declare no potential conflicts of interest with respect to the research, authorship, and/or publication of this article.

