## Supplemental Figures and Tables for "*brca2*-mutant zebrafish exhibit context- and tissue-dependent alterations in cell phenotypes and response to injury"

**Supplementary Information**

1. Supplementary Figure Legends
2. Supplementary Figures S1-S3
3. Supplementary Tables S1-S4

**Supplementary Figure S1. RNA probe and antibody validations and negative controls for *in situ* hybridization and immunohistochemistry experiments.** Areas boxed in lettered figures are shown to the right. **(a)** *blbp* expression in the brain (red chromogen). **(b)** sox2 expression in the brain (brown chromogen). Red and yellow arrows indicate positive and negative nuclei, respectively. Yellow asterisk, nucleated red blood cells within the choroid rete (negative for sox2 expression). **(c)** sox10 expression in the brain (brown chromogen). Red and yellow arrows indicate positive and negative nuclei, respectively. Yellow asterisk, nucleated red blood cells within the choroid rete (negative for sox10 expression). **(d)** lcp1 expression in the spleen (purple chromogen). **(e)** Negative control probe for RNA *in situ* hybridization experiments (red chromogen). **(f)** Secondary-only antibody control with hematoxylin counterstain for sox2 and sox10 immunohistochemistry experiments (brown chromogen). **(g)** Secondary-only antibody control with methyl green counterstain for lcp1 and krt18 immunohistochemistry experiments (purple chromogen).

**Supplementary Figure S2. Quantitative analysis of sox2-, sox10-, and krt18-expressing cells in the zebrafish optic nerve after unilateral optic nerve injury. (a)** Representative examples of sox2 expression (brown chromogen), a marker for neural stem cells, in the injured and uninjured optic nerve. Red and yellow arrows indicate positive and negative nuclei, respectively. **(b,c)** Quantitative analysis of sox2-positive cells in the optic nerve three days (b) and two weeks (c) post-injury. **(d)** Examples of sox10 expression (brown chromogen), a marker for neural crest progenitor cells, oligodendrocytes, and Schwann cell precursors, in the injured and uninjured optic nerve. Red and yellow arrows indicate positive and negative nuclei, respectively. **(e,f)** Quantitative analysis of sox10-positive cells in the optic nerve three days (e) and two weeks (f) post-injury. **(g)** Representative examples of krt18 expression (purple chromogen), a marker for reactive astrocytes in zebrafish optic nerve, in the injured and uninjured optic nerve. **(h,i)** Quantitative analysis of krt-18 positive area in the uninjured versus injured optic nerve three days (h) and two weeks (i) post-injury. Ret, retina; CR, choroid rete; ON, optic nerve, UI, uninjured; I, injured. Scale bar = 100 μm (panels a, d); 50 μm (panel g).

**Supplementary Figure S3. Semi-quantitative analysis of *blbp* and sox2 expression in zebrafish ocular tumors**. Expression of *blbp* (red chromogen) and sox2 (brown chromogen) is low or absent in the majority of ocular tumors regardless of optic nerve injury (ONI) status. Tumor invasion and subsequent optic nerve/retinal disruption induces marked expression of *blbp* in the retina. Arrows indicated dispersed melanin pigment within the tumor derived from the disrupted RPE. Ret, retina; RPE, retinal pigmented epithelium. Scale bar = 50 μm.

**Supplementary Figure S1**

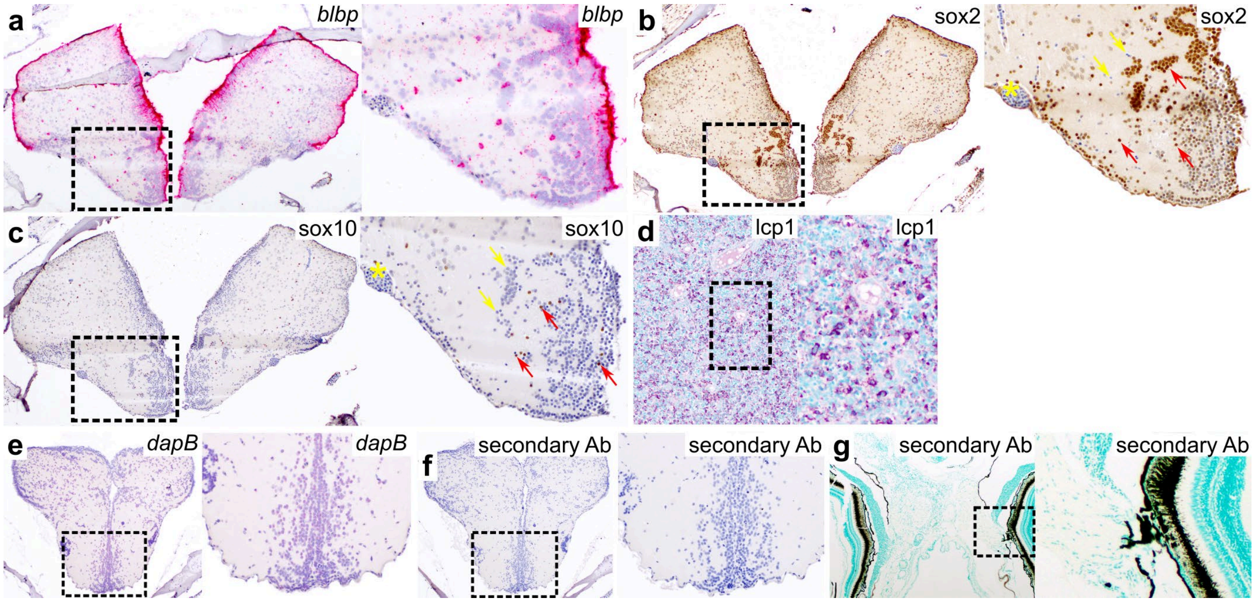

**Supplementary Figure S2**

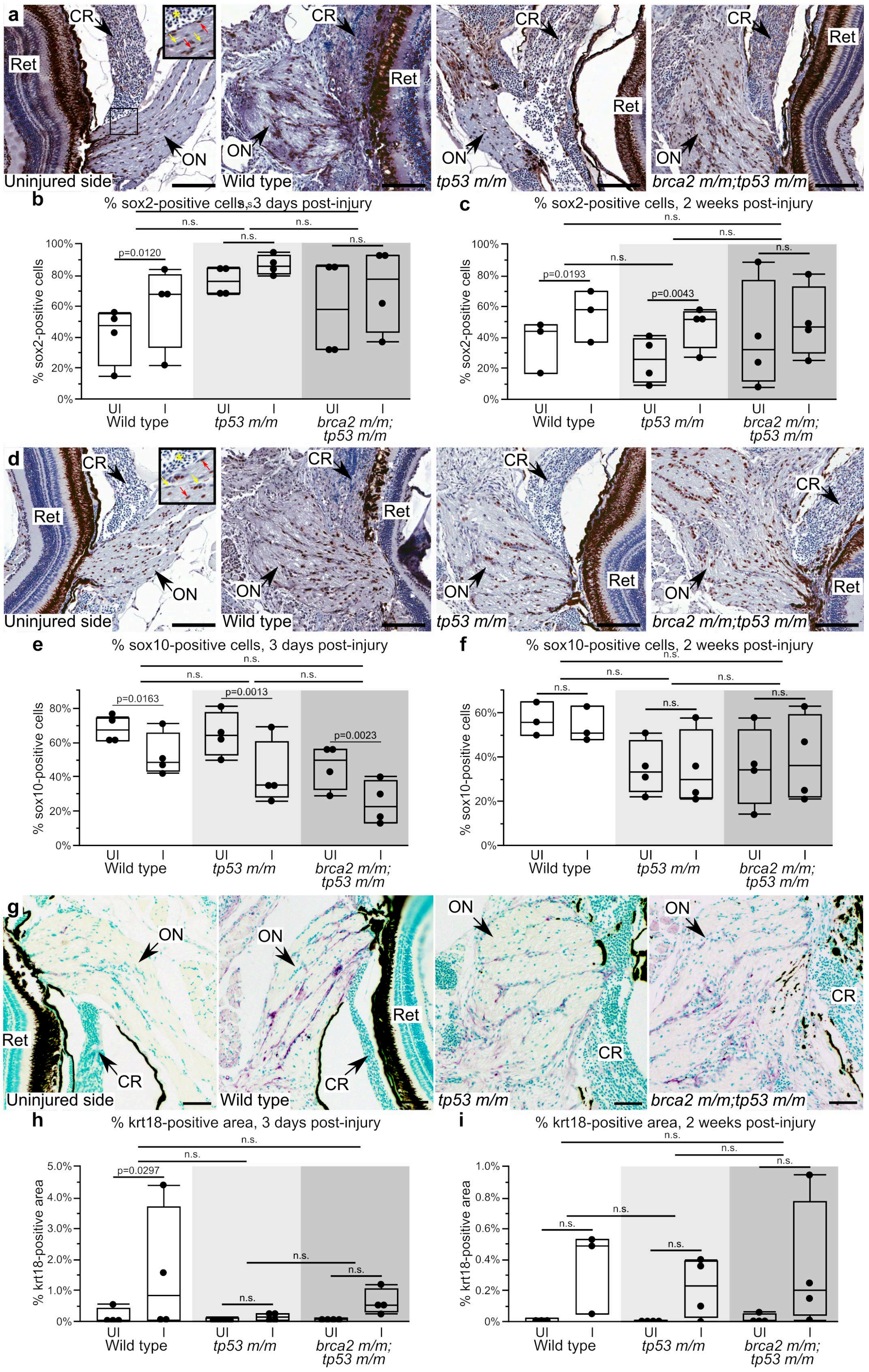

### Supplementary Figure S3

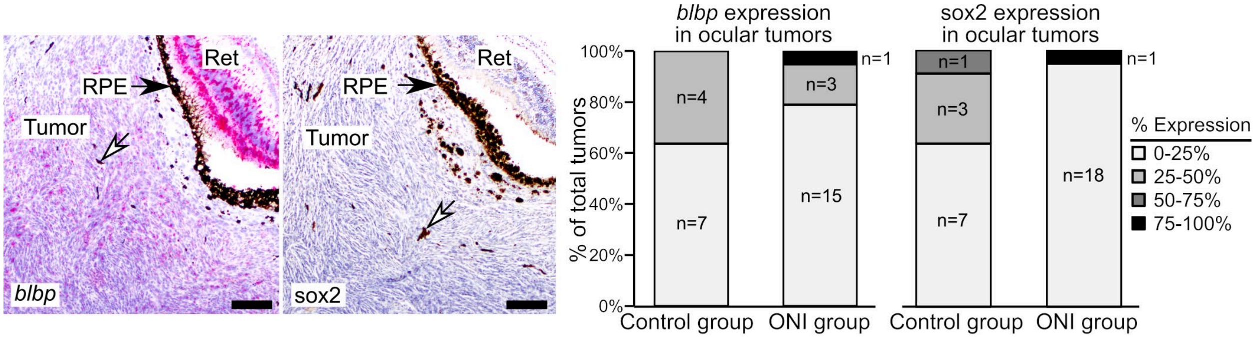

**Supplementary Table S1. Statistical comparisons used to assess the relationships between genotype, optic nerve injury (ONI) status, and phenotype of ocular lesions.** CR, choroid rete; mo, months. Statistically significant p-values (p<0.05) are in bold type.

| | **Statistical test** | **p-value** |
|---|---|---|
| **Egg/embryo phenotype** | | |
| Day 0, *tp53 m/m* versus *brca2 m/m;tp53 m/m* | Fisher's Exact Test | **<0.0001** |
| Day 1, *tp53 m/m* versus *brca2 m/m;tp53 m/m* | Fisher's Exact Test | **<0.0001** |
| **Site of tumor development (ocular or non-ocular)** | | |
| ONI versus control | Fisher's Exact Test | p=0.6789 |
| *tp53 m/m* ONI versus control | Fisher's Exact Test | p=0.6680 |
| *brca2 m/m;tp53 m/m* ONI versus control | Fisher's Exact Test | p=1.0000 |
| **Side of ocular tumor development (right/ONI, left/non-ONI, or bilateral)** | | |
| ONI versus control | Fisher's Exact Test | p=0.8328 |
| *tp53 m/m* ONI versus control | Fisher's Exact Test | p=0.6837 |
| *brca2 m/m;tp53 m/m* ONI versus control | Fisher's Exact Test | p=0.8679 |
| **Presence of atypical spindle cells in the CR of zebrafish without ocular tumors**[a] | | |
| *tp53 m/m* versus *brca2 m/m;tp53 m/m* | Fisher's Exact Test | p=0.1941 |
| **Presence of ocular lesion (ocular tumor, CR spindle cells) in zebrafish after ONI**[a] | | |
| *brca2 m/m;tp53 m/m* versus *tp53 m/m* | Fisher's Exact Test | **p=0.0220** |
| **Bilateral or unilateral ocular lesion in zebrafish after ONI**[b] | | |
| *brca2 m/m;tp53 m/m* versus *tp53 m/m* | Fisher's Exact Test | p=0.1207 |

[a] 4 zebrafish (*tp53 m/m*, n=1; *brca2 m/m;tp53 m/m*, n=3) were excluded from this comparison because the choroid rete could not be evaluated sufficiently on both sides in histologic sections.
[b] 5 zebrafish (*tp53 m/m*, n=2; *brca2 m/m;tp53 m/m*, n=3) were excluded from this comparison because the choroid rete could not be evaluated sufficiently on one side in histologic sections.

**Supplementary Table S2. Statistical comparisons used to assess the relationship between optic nerve injury (ONI) status, genotype, and characteristics of the cell population in the optic nerve.** I, injured; UI, uninjured. Statistically significant p-values ($p<0.05$) are in bold type.

| Comparison | p-value | Difference and confidence interval |
|---|---|---|
| ***Total cellularity, three days post-injury*** | | |
| Wild type (I vs. UI) | 0.9554 | Difference, -0.01; Lower, -0.46; Upper, 0.43 |
| *tp53 m/m* (I vs. UI) | 0.6805 | Difference, 0.09; Lower, -0.35; Upper, 0.53 |
| *brca2 m/m;tp53 m/m* (I vs. UI) | **0.0001** | Difference, 0.95; Lower, 0.56; Upper, 1.34 |
| Wild type (I-UI) vs. *tp53 m/m* (I-UI) | 0.7408 | Difference, -0.10; Lower, -0.73; Upper, 0.53 |
| Wild type (I-UI) vs. *brca2 m/m;tp53 m/m* (I-UI) | **0.0028** | Difference, -0.96; Lower, -1.55; Upper, -0.37 |
| *tp53 m/m* (I-UI) vs. *brca2 m/m;tp53 m/m* (I-UI) | **0.0064** | Difference, -0.86; Lower, -1.45; Upper, -0.27 |
| ***Total cellularity, two weeks post-injury*** | | |
| Wild type (I vs. UI) | **0.0464** | Difference, 0.69; Lower, 0.01; Upper, 1.36 |
| *tp53 m/m* (I vs. UI) | **0.0130** | Difference, 0.91; Lower, 0.24; Upper, 1.58 |
| *brca2 m/m;tp53 m/m* (I vs. UI) | **0.0000** | Difference, 1.83; Lower, 1.23; Upper, 2.43 |
| Wild type (I-UI) vs. *tp53 m/m* (I-UI) | 0.6107 | Difference, -0.22; Lower, -1.18; Upper, 0.73 |
| Wild type (I-UI) vs. *brca2 m/m;tp53 m/m* (I-UI) | **0.0179** | Difference, -1.15; Lower, -2.05; Upper, -0.24 |
| *tp53 m/m* (I-UI) vs. *brca2 m/m;tp53 m/m* (I-UI) | **0.0463** | Difference, -0.92; Lower, -1.82; Upper, -0.02 |
| ***blbp expression, three days post-injury*** | | |
| Wild type (I vs. UI) | **0.0039** | Difference, 0.05; Lower, 0.02; Upper, 0.09 |
| *tp53 m/m* (I vs. UI) | **0.0179** | Difference, 0.04; Lower, 0.01; Upper, 0.07 |
| *brca2 m/m;tp53 m/m* (I vs. UI) | **0.0001** | Difference, 0.09; Lower, 0.06; Upper, 0.13 |
| Wild type (I-UI) vs. *tp53 m/m* (I-UI) | 0.5134 | Difference, 0.01; Lower, -0.03; Upper, 0.06 |
| Wild type (I-UI) vs. *brca2 m/m;tp53 m/m* (I-UI) | 0.0757 | Difference, -0.04; Lower, -0.08; Upper, 0.01 |
| *tp53 m/m* (I-UI) vs. *brca2 m/m;tp53 m/m* (I-UI) | **0.0249** | Difference, -0.05; Lower, -0.10; Upper, -0.01 |
| ***blbp expression, two weeks post-injury*** | | |
| Wild type (I vs. UI) | 0.9207 | Difference, 0.00; Lower, -0.08; Upper, 0.09 |
| *tp53 m/m* (I vs. UI) | 0.7072 | Difference, 0.01; Lower, -0.06; Upper, 0.08 |
| *brca2 m/m;tp53 m/m* (I vs. UI) | **0.0071** | Difference, 0.11; Lower, 0.04; Upper, 0.18 |
| Wild type (I-UI) vs. *tp53 m/m* (I-UI) | 0.8637 | Difference, -0.01; Lower, -0.12; Upper, 0.10 |
| Wild type (I-UI) vs. *brca2 m/m;tp53 m/m* (I-UI) | 0.0526 | Difference, -0.11; Lower, -0.22; Upper, 0.00 |
| *tp53 m/m* (I-UI) vs. *brca2 m/m;tp53 m/m* (I-UI) | 0.0533 | Difference, -0.10; Lower, -0.20; Upper, 0.00 |
| ***lcp1 expression, three days post-injury*** | | |
| Wild type (I vs. UI) | 0.7887 | Difference, 0.00; Lower, 0.00; Upper, 0.00 |
| *tp53 m/m* (I vs. UI) | 0.1904 | Difference, 0.00; Lower, 0.00; Upper, 0.00 |
| *brca2 m/m;tp53 m/m* (I vs. UI) | **0.0374** | Difference, 0.00; Lower, 0.00; Upper, 0.01 |
| Wild type (I-UI) vs. *tp53 m/m* (I-UI) | 0.4409 | Difference, 0.00; Lower, -0.01; Upper, 0.00 |
| Wild type (I-UI) vs. *brca2 m/m;tp53 m/m* (I-UI) | 0.1605 | Difference, 0.00; Lower, -0.01; Upper, 0.00 |
| *tp53 m/m* (I-UI) vs. *brca2 m/m;tp53 m/m* (I-UI) | 0.4878 | Difference, 0.00; Lower, 0.00; Upper, 0.00 |
| ***lcp1 expression, two weeks post-injury*** | | |
| Wild type (I vs. UI) | 0.9935 | Difference, 0.00; Lower, -0.01; Upper, 0.01 |
| *tp53 m/m* (I vs. UI) | 0.8692 | Difference, 0.00; Lower, -0.01; Upper, 0.01 |
| *brca2 m/m;tp53 m/m* (I vs. UI) | **0.0372** | Difference, 0.01; Lower, 0.00; Upper, 0.02 |
| Wild type (I-UI) vs. *tp53 m/m* (I-UI) | 0.9190 | Difference, 0.00; Lower, -0.01; Upper, 0.01 |
| Wild type (I-UI) vs. *brca2 m/m;tp53 m/m* (I-UI) | 0.1422 | Difference, -0.01; Lower, -0.02; Upper, 0.00 |
| *tp53 m/m* (I-UI) vs. *brca2 m/m;tp53 m/m* (I-UI) | 0.1386 | Difference, -0.01; Lower, -0.02; Upper, 0.00 |

**Supplementary Table S3. Statistical comparisons used to assess the relationship between optic nerve injury (ONI) status, genotype, and characteristics of the cell population in the optic nerve.** I, injured; UI, uninjured. Statistically significant p-values ($p<0.05$) are in bold type.

| Comparison | p-value | Difference and confidence interval |
|---|---|---|
| ***sox2 expression, three days post-injury*** | | |
| Wild type (I vs. UI) | **0.0120** | Difference, 0.19; Lower, 0.05; Upper, 0.33 |
| *tp53 m/m* (I vs. UI) | 0.1205 | Difference, 0.10; Lower, -0.03; Upper, 0.24 |
| *brca2 m/m;tp53 m/m* (I vs. UI) | 0.0708 | Difference, 0.12; Lower, -0.01; Upper, 0.26 |
| Wild type (I-UI) vs. *tp53 m/m* (I-UI) | 0.3416 | Difference, 0.09; Lower, -0.11; Upper, 0.28 |
| Wild type (I-UI) vs. *brca2 m/m;tp53 m/m* (I-UI) | 0.4621 | Difference, 0.07; Lower, -0.13; Upper, 0.26 |
| *tp53 m/m* (I-UI) vs. *brca2 m/m;tp53 m/m* (I-UI) | 0.8187 | Difference, -0.02; Lower, -0.21; Upper, 0.17 |
| ***sox2 expression, two weeks post-injury*** | | |
| Wild type (I vs. UI) | **0.0193** | Difference, 0.19; Lower, 0.04; Upper, 0.34 |
| *tp53 m/m* (I vs. UI) | **0.0043** | Difference, 0.22; Lower, 0.09; Upper, 0.35 |
| *brca2 m/m;tp53 m/m* (I vs. UI) | 0.1262 | Difference, 0.10; Lower, -0.03; Upper, 0.22 |
| Wild type (I-UI) vs. *tp53 m/m* (I-UI) | 0.7163 | Difference, -0.03; Lower, -0.23; Upper, 0.16 |
| Wild type (I-UI) vs. *brca2 m/m;tp53 m/m* (I-UI) | 0.3079 | Difference, 0.09; Lower, -0.10; Upper, 0.29 |
| *tp53 m/m* (I-UI) vs. *brca2 m/m;tp53 m/m* (I-UI) | 0.1521 | Difference, 0.12; Lower, -0.06; Upper, 0.31 |
| ***sox10 expression, three days post-injury*** | | |
| Wild type (I vs. UI) | **0.0163** | Difference, -0.15; Lower, -0.27; Upper, -0.04 |
| *tp53 m/m* (I vs. UI) | **0.0013** | Difference, -0.24; Lower, -0.35; Upper, -0.12 |
| *brca2 m/m;tp53 m/m* (I vs. UI) | **0.0023** | Difference, -0.22; Lower, -0.33; Upper, -0.10 |
| Wild type (I-UI) vs. *tp53 m/m* (I-UI) | 0.2688 | Difference, 0.09; Lower, -0.08; Upper, 0.25 |
| Wild type (I-UI) vs. *brca2 m/m;tp53 m/m* (I-UI) | 0.3986 | Difference, 0.06; Lower, -0.10; Upper, 0.23 |
| *tp53 m/m* (I-UI) vs. *brca2 m/m;tp53 m/m* (I-UI) | 0.7767 | Difference, -0.02; Lower, -0.19; Upper, 0.14 |
| ***sox10 expression, two weeks post-injury*** | | |
| Wild type (I vs. UI) | 0.6868 | Difference, -0.03; Lower, -0.19; Upper, 0.13 |
| *tp53 m/m* (I vs. UI) | 0.9998 | Difference, 0.00; Lower, -0.14; Upper, 0.14 |
| *brca2 m/m;tp53 m/m* (I vs. UI) | 0.5498 | Difference, 0.04; Lower, -0.10; Upper, 0.17 |
| Wild type (I-UI) vs. *tp53 m/m* (I-UI) | 0.7601 | Difference, -0.03; Lower, -0.24; Upper, 0.18 |
| Wild type (I-UI) vs. *brca2 m/m;tp53 m/m* (I-UI) | 0.4892 | Difference, -0.07; Lower, -0.27; Upper, 0.14 |
| *tp53 m/m* (I-UI) vs. *brca2 m/m;tp53 m/m* (I-UI) | 0.6705 | Difference, -0.04; Lower, -0.23; Upper, 0.16 |
| ***krt18 expression, three days post-injury*** | | |
| Wild type (I vs. UI) | **0.0297** | Difference, 0.01; Lower, 0.00; Upper, 0.03 |
| *tp53 m/m* (I vs. UI) | 0.8537 | Difference, 0.00; Lower, -0.01; Upper, 0.01 |
| *brca2 m/m;tp53 m/m* (I vs. UI) | 0.3186 | Difference, 0.01; Lower, -0.01; Upper, 0.02 |
| Wild type (I-UI) vs. *tp53 m/m* (I-UI) | 0.1253 | Difference, 0.01; Lower, 0.00; Upper, 0.03 |
| Wild type (I-UI) vs. *brca2 m/m;tp53 m/m* (I-UI) | 0.3092 | Difference, 0.01; Lower, -0.01; Upper, 0.03 |
| *tp53 m/m* (I-UI) vs. *brca2 m/m;tp53 m/m* (I-UI) | 0.5554 | Difference, 0.00; Lower, -0.02; Upper, 0.01 |
| ***krt18 expression, two weeks post-injury*** | | |
| Wild type (I vs. UI) | 0.0746 | Difference, 0.00; Lower, 0.00; Upper, 0.01 |
| *tp53 m/m* (I vs. UI) | 0.1891 | Difference, 0.00; Lower, 0.00; Upper, 0.01 |
| *brca2 m/m;tp53 m/m* (I vs. UI) | 0.0636 | Difference, 0.00; Lower, 0.00; Upper, 0.01 |
| Wild type (I-UI) vs. *tp53 m/m* (I-UI) | 0.5591 | Difference, 0.00; Lower, 0.00; Upper, 0.01 |
| Wild type (I-UI) vs. *brca2 m/m;tp53 m/m* (I-UI) | 0.8916 | Difference, 0.00; Lower, 0.00; Upper, 0.01 |
| *tp53 m/m* (I-UI) vs. *brca2 m/m;tp53 m/m* (I-UI) | 0.6262 | Difference, 0.00; Lower, -0.01; Upper, 0.00 |

**Supplementary Table S4. Experimental groups**. ONI, optic nerve injury; dpi, days post-injury; wpi, weeks post-injury; IHC, immunohistochemistry; ISH, in situ hybridization; mo, months.

| **Evaluation of ovaries** | |
|---|---|
| **Genotype** | **7.5 mo** |
| Wild type | 4 |
| *tp53 m/m* | 4 |
| *brca2 m/m;tp53 m/m* | 4 |

| **Serial sacrifice group** | | | | | | |
|---|---|---|---|---|---|---|
| **Genotype** | **3 mo** | **4 mo** | **5.1 mo**[a] | **5.5 mo**[a] | **6 mo** | **7 mo** |
| Wild type | -- | -- | 4 | 4 | -- | 4 |
| *tp53 m/m* | 4 | 4 | 4 | 4 | 4 | 4 |
| *brca2 m/m;tp53 m/m* | 4 | 4 | 4 | 4 | 4 | 4 |

| **ONI group for short-term injury response** | | |
|---|---|---|
| **Genotype** | **3 dpi (ONI at 5 mo)** | **2 wpi (ONI at 5 mo)** |
| Wild type | 7 | 4 |
| *tp53 m/m* | 7 | 4 |
| *brca2 m/m;tp53 m/m* | 9 | 5 |

| **ONI group for tumorigenesis studies** | | |
|---|---|---|
| **Genotype** | **Total zebrafish**[a] | **Zebrafish with tumors**[b] |
| Wild type | 36 | 0 (0%) |
| *tp53 m/m* | 54 | 47 (87%) |
| *brca2 m/m;tp53 m/m* | 60 | 56 (93%) |

| **Control group for tumorigenesis studies** | | |
|---|---|---|
| **Genotype** | **Total zebrafish**[c,d] | **Zebrafish with tumors**[e] |
| *tp53 m/m* | 55 | 48 (87%) |
| *brca2 m/m;tp53 m/m* | 49 | 46 (94%) |

| **ONI group for IHC and ISH analysis of ocular tumors** | | | | |
|---|---|---|---|---|
| **Genotype** | **Total tumors** | **Right (ONI side)** | **Left (non-ONI side)** | **Bilateral** |
| *tp53 m/m* | 7 | 3 | 3 | 1 |
| *brca2 m/m;tp53 m/m* | 12 | 5 | 3 | 4 |

| **Control group for IHC and ISH analysis of ocular tumors**[f] | | | | |
|---|---|---|---|---|
| **Genotype** | **Total tumors** | **Right (ONI side)** | **Left (non-ONI side)** | **Bilateral** |
| *tp53 m/m* | 5 | 2 | 3 | 0 |
| *brca2 m/m;tp53 m/m* | 6 | 2 | 4 | 0 |

[a] 8 zebrafish (*tp53 m/m*, n=3; *brca2 m/m;tp53 m/m*, n=5) were excluded from this group (see Methods)
[b] 18 zebrafish (*tp53 m/m*, n=6; *brca2 m/m;tp53 m/m*, n=12) developed two anatomically distinct tumors (e.g, an ocular tumor and a non-ocular tumor). In these cases, tumors were counted separately.
[c] The control group is a single clutch of zebrafish siblings that was previously described in an analysis of tumor ploidy, and includes all tumors identified in this population. Not all tumors from this population were included in the previous study (e.g, excluded because ploidy analysis did not pass QC analysis)[27].
[d] 11 zebrafish (*tp53 m/m*, n=6; *brca2 m/m;tp53 m/m*, n = 5) were excluded from this group (see Methods)
[e] 12 zebrafish (*tp53 m/m*, n=3; *brca2 m/m;tp53 m/m*, n=9) developed two anatomically distinct tumors (e.g, an ocular tumor and a non-ocular tumor). In these cases, tumors were counted separately.
[f] Ocular tumors were derived from a population of zebrafish with *brca2* and *tp53* mutations that were collected and processed at the same time as the ONI group in order to minimize differences in tissue orientation, fixation, processing, and specimen age.
